# TRAIT: A Comprehensive Database for T-cell Receptor-Antigen Interactions

**DOI:** 10.1101/2024.11.20.624436

**Authors:** Mengmeng Wei, Jingcheng Wu, Shengzuo Bai, Yuxuan Zhou, Yichang Chen, Xue Zhang, Wenyi Zhao, Ying Chi, Gang Pan, Feng Zhu, Shuqing Chen, Zhan Zhou

## Abstract

Comprehensive and integrated resources on interactions between T-cell receptors and antigens are still lacking for adoptive T-cell-based immunotherapies, highlighting a significant gap that must be addressed to fully comprehend the mechanisms of antigen recognition by T-cells. In this study, we present TRAIT, a comprehensive database that profiles the interactions between T-cell receptors (TCRs) and antigens. TRAIT stands out due to its comprehensive description of TCR-antigen interactions by integrating sequences, structures and affinities. It provides nearly 8 million experimentally validated TCR-antigen pairs, resulting in an exhaustive landscape of antigen-specific TCRs. Notably, TRAIT emphasizes single-cell omics as a major reliable data source for TCR-antigen interactions and includes millions of reliable non-interactive TCRs. Additionally, it thoroughly demonstrates the interactions between mutations of TCRs and antigens, thereby benefiting affinity maturation of engineered TCRs as well as vaccine design. TCRs on clinical trials were innovatively provided. With the significant efforts made towards elucidating the complex interactions between TCRs and antigens, TRAIT is expected to ultimately contribute superior algorithms and substantial advancements in the field of T-cell-based immunotherapies. TRAIT is freely accessible at https://pgx.zju.edu.cn/traitdb.

## Introduction

Remarkable T-cell receptors (TCRs) play a vital role in initiating intracellular signals required for the effector response to foreign antigens in the adaptive immune system [1,2]. TCR provides the initial signal for T-cell activation by mediating the specific recognition of pathogen-derived epitopes through its interaction with the peptide-major histocompatibility complex (pMHC) multimer [3–5]. The underlying basis for the exceptional sensitivity and discrimination recognition lies in the hypervariable complementarity-determining region 3 (CDR3) loops of TCRs, which interact mainly with the peptide, whereas the CDR1 and CDR2 regions contact the MHC α-helices [6,7]. The vast diversity of CDR3 regions and the cross-reactivity of TCRs ensure that the available TCR repertoire can recognize a wide range of pathogen-derived epitopes encountered over a lifetime. Nevertheless, the rules of how TCRs recognize cognate antigens (pMHCs) remain poorly understood [8,9]. Therefore, a precise and thorough characterization of the interactions between TCRs and antigens is important for understanding the molecular mechanisms of pathogen elimination and tumor surveillance mediated by adaptive immune responses.

Owing to their great importance, data including sequences, structures and specificities of TCRs, TCR repertoires from different donors, and affinities of TCR-antigen pairs and their mutations have attracted extensive interest [10–12]. Furthermore, these data are essential for the fields of TCR engineering and the development of immunotherapies [13–15]. To date, several TCR-related databases have been developed and are currently active [16–26], the majority of which focus on providing structural data of TCRs or TCR-pMHC complexes (*e.g.*, STCRDab [16], TCR3d [17], and ATLAS [18]), and others are specialized in describing sequences of TCR repertoires (e.g., UcTCRdb [19], PIRD [20], and TCRdb [21]). Additionally, several reputable databases (e.g., McPAS-TCR [22], VDJdb [23,24], neoTCR [25], and IEDB [26]) have demonstrated a wealth of information on TCRs with known antigen specificity. Although databases containing valuable TCR data are available, most existing ones concentrate on a specific aspect. Binding affinity and clinical evaluation of TCRs are not typically included in the majority of active databases (**Table 1**). There is noticeable gap when it comes to an integrated database that compiles a landscape of TCR-antigen interactions, highlighting the urgent need for a comprehensive database profiling TCR-antigen interactions.

**Table 1.** An overview of databases related to TCRs.

| Database | Sequences | Structures | Specificities | Binding<br>affinity | Non-interactive<br>pairs | Mutations | Clinical<br>evaluation | Active | Reference |
| --- | --- | --- | --- | --- | --- | --- | --- | --- | --- |
| STCRDab |  | ✓ |  |  |  |  |  | ✓ | [16] |
| TCR3d |  | ✓ |  |  |  |  |  | ✓ | [17] |
| ATLAS |  | ✓ |  | ✓ |  | ✓ |  | No longer updated after 2017 | [18] |
| TBAdb | ✓ |  |  |  |  |  |  | ✓ | [20] |
| TCRdb | ✓ |  |  |  |  |  |  | ✓ | [21] |
| McPAS-TCR | ✓ |  | ✓ |  |  |  |  | ✓ | [22] |
| VDJdb | ✓ | ✓ | ✓ |  |  |  |  | ✓ | [23] |
| NeoTCR | ✓ |  | ✓ |  |  |  |  | ✓ | [25] |
| IEDB | ✓ |  | ✓ |  |  |  |  | ✓ | [26] |
| TRAIT | ✓ | ✓ | ✓ | ✓ | ✓ | ✓ | ✓ | ✓ | This paper |

In this study, we developed a novel knowledge base focused on the T-cell receptor-antigen interactions (TRAIT). First, an extensive literature review was conducted by performing keyword searches in PubMed, public datasets and omics platform, resulting in a total of 7,979,361 antigen-specificity validated TCR sequences for 1,127 unique epitopes and 112 MHC alleles. Uniquely, 7,927,923 non-interactive TCR sequences verified by single-cell omics were collected and recorded. Secondly, structures of TCR-antigen complexes were then systematically collected from the Protein Data Bank (PDB) [27], and a total of 220 structures were identified. And then, valuable 3D affinity data quantifying the binding ability of TCRs and antigens were manually retrieved, leading to 759 binding affinity records. A unique compilation consisting of 564 TCR mutants and 467 antigen mutants were collected and associated with their respective binding affinities. Finally, TCRs with clinical evaluations were consulted exhaustively and 34 clinical trials inferring 65 antigen-specific TCRs were identified.

In summary, TRAIT was developed to explicitly characterize the interactions between TCRs and antigens by integrating sequences, structures and binding affinities. TRAIT also offers single-cell-based antigen-specific TCR repertoires and systematically demonstrates interactions between mutations of TCRs and antigens. Additionally, clinical evaluations of antigen-specific TCRs were uniquely provided. Given the recent prominence of T-cell-based immunotherapies in disease treatment, the comprehensive data on TCR-antigen interactions provided by TRAIT are anticipated to better facilitate the discovery and optimization of TCR candidates for immunotherapies.

## Data collection and processing

### Collection and processing of interaction data between TCRs and antigens

Based on literature evidence of TCR-antigen pairs including sequences, structures and binding affinities, were first collected by searching PubMed [28] for keywords with the combined terms ‘T cell receptor + repertoire’, ‘T cell receptor + affinity’, ‘T cell receptor + specificity’, ‘T cell receptor + major histocompatibility complexes’, and so on. Sequences of antigen-specific TCRs were also collected from public databases such as VDJdb [24], McPAS-TCR [22], IEDB [26], and omics datasets available on the 10X website. The available raw data were downloaded and processed through IMGT/V-QUEST to produce annotated information for TCRs that lacked V and/or J specifications or had incomplete or overly extended CDR3 sequences. Structures of TCR-antigen complexes were further identified from PDB [27] with TCR-antigen reference sequences as queries. Only structures of TCR-MHC or TCR-pMHC complexes were collected and associated with corresponding sequence entries in TRAIT. The binding affinities of TCR-antigen pairs experimentally determined using purified proteins by surface plasmon resonance (SPR) or isothermal titration calorimetry (ITC) were collected and recorded. Finally, the manually collected data of TCR-antigen pairs were further checked to remove duplicates.

### Single-cell based antigen-specific TCRs identification

Datasets on the 10X website that describe TCRs extracted from pMHC multimer-labeled T-cells and identified by single-cell sequencing were collected and recorded as ‘interactive pairs’ in TRAIT. Simultaneously, TCRs extracted from non-binding T-cell populations after staining with a pMHC multimer and identified by single-cell sequencing were collected and recorded as ‘non-interactive pairs’.

### Mutations classification and referring to binding affinity

A systemic literature review was conducted using combinations of keywords such as ‘T cell receptors + mutations’, ‘T cell receptors + variants’, ‘T cell receptors + directed evolution’, ‘T cell receptors + affinity maturation’, ‘major histocompatibility complex + mutations’, ‘major histocompatibility complex + variants’, and so on. Mutations of TCRs or antigens with binding affinities measured experimentally were collected and recorded in TRAIT. All of the entries were divided into two categories including ‘mutations of TCRs’ and ‘mutations of antigens’ according to the sites of mutated amino acids. CDR sequences of TCRs were annotated through IMGT/V-QUEST by inputting their full-length sequences.

### Collection of TCRs on clinical trials

A thorough literature review in diverse was conducted across multiple official patent organizations, various pharmaceutical company pipelines, and literature databases using keywords ‘TCR-T’, ‘TCR-transduced T cell’, ‘TCR-engineered T cell’, and so on. Detailed activity data of TCR gene-engineered T cells (TCR-T) including objective response rates (ORRs), clinical response, and toxicities related to TCR-T in different stages of clinical trials were collected and displayed in TRAIT.

### Database contents and construction

As a result, nearly 8 million records of TCR/antigen pairs associated with infectious diseases (including COVID-19), cancer and autoimmune diseases were collected, and the statistical data are presented in **Table 2**. Large-scale TCR/antigen pairs including sequences (7,979,361 records), structures (220 records) and binding affinities (759 records) were manually retrieved. Multiple antigen species, including *Homo Sapiens* (3,091), SARS-CoV-2 (4,072), Influenza A (9,862), EBV (6,230), CMV (22,295), and HIV (2,656), were involved, and a total of 112 MHC alleles were extracted. Ultimately, a total of 20,251 interactive TCRs and 7,927,923 non-interactive TCRs referring to 50 antigens from four different donors were collected and recorded in TRAIT. And totally 1031 mutation records, including 564 TCR mutations referring to 29 wild-type antigens and 467 antigen mutants referring to 25 wild-type TCRs were presented. Additionally, TRAIT innovatively compiles information on 34 TCR-T cell therapies that are in clinical trial stage and have published their clinical outcomes. 65 TCR-antigen pairs (59 wild-type TCRs and 6 modified TCRs) and 25 cancer types were inferred.

**Table 2.** Statistics recorded by epitope species in TRAIT.

| Epitope species | Unique epitopes | Interactive TCR | Non-interactive TCR | Structures | Affinities | Mutations of TCRs | Mutations of antigens |
| --- | --- | --- | --- | --- | --- | --- | --- |
| *Homo Sapiens* | 217 | 3,103 | 3,200,768 | 81 | 20 | 168 | 61 |
| Influenza A | 22 | 9,862 | 151,171 | 16 | 153 | 23 | 4 |
| SARS-CoV-2 | 668 | 4,072 | 6 | 4 | 5 | 0 | 0 |
| EBV | 30 | 6,230 | 1,887,219 | 15 | 50 | 65 | 53 |
| CMV | 31 | 22,295 | 1,408,426 | 6 | 233 | 2 | 0 |
| HIV | 53 | 2,656 | 800,204 | 23 | 128 | 28 | 9 |
| HPV | 1 | 12 | 320,104 | 0 | 30 | 0 | 0 |
| HCV | 15 | 1,111 | 0 | 5 | 0 | 0 | 6 |
| HTLV-1 | 8 | 178 | 160,013 | 9 | 0 | 114 | 20 |
| Others | 82 | 1,919 | 12 | 61 | 140 | 164 | 314 |
| Total | 1,127 | 51,438 | 7,927,923 | 220 | 759 | 564 | 467 |

*Note:* SARS-CoV-2, severe acute respiratory syndrome coronavirus 2; EBV, Epstein-Barr virus; CMV, cytomegalovirus; HIV, human immunodeficiency virus; HPV, human papilloma virus; HCV, hepatitis C virus; HTLV-1, human T-lymphotropic virus type 1.

TRAIT was built using PHP (https://www.php.net), Bootstrap (https://getbootstrap.com), and WordPress (https://wordpress.org), a PHP-based blogging platform that enables users to create websites on servers supporting PHP and MySQL databases. JQuery (https://jquery.com) and DataTables are employed to showcase the query result tables. All data in this database is stored and managed in MySQL. And the workflow of TRAIT is depicted in **Figure 1**.

**Figure 1.**
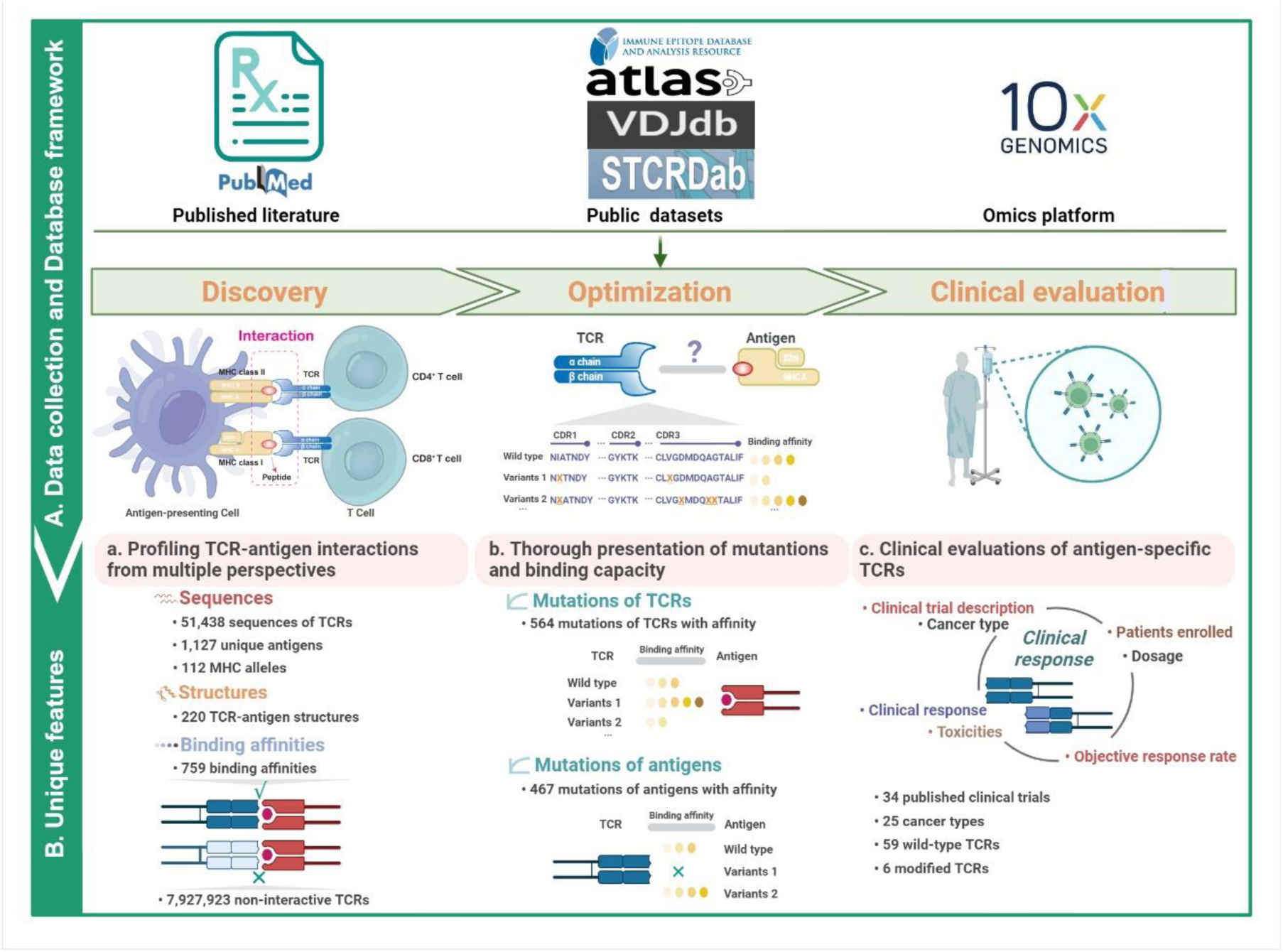
Overview of the TRAIT database. **A**. Data collection and database framework. Information of antigen-specific TCRs including sequences, structures, affinities, mutations with binding capacities and clinical response, were thoroughly collected from published literatures, public databases and omics platform. And then data were reorganized by the whole process of ‘discovery-optimization-clinical evaluations’ of antigen-specific TCRs. **B**. Unique features of TRAIT. TRAIT is unique in (a) unambiguously profiling the interactions between T-cell receptors and antigens by integrating sequences, structures, and affinities, thus offering an exhaustively landscape of TCR-antigen interactions, (b) systematically describing mutations-affinity relationship of TCRs and antigens, benefiting affinity maturation of TCRs, and **(c)** thoroughly demonstrating clinical response of TCRs on clinical trials. The figure is created with BioRender.com.

### User interface

TRAIT has a user-friendly web interface containing seven pages: ‘Home’, ‘Search’, ‘Omics’, ‘Mutations’, ‘Therapeutics’, ‘Download’ and ‘Help’ (see **Figure 2**). The ‘Home’ page provides an introduction to the entire database, an overview of collected data, and a visualization of total records. The ‘Search’ page provides a comprehensive repository of TCR-antigen pairs in TRAIT. The query term can be either a CDR3 sequence of TCRs (Nucleic acid or amino acid sequences), information of antigens (epitope, parent gene, species, the MHC allele), interactive information (binding affinity, structure), or metadata (PMID of reference, sequencing method of TCR, publication year). A built-in search box allows users to refine results to a specific subject by entering additional search terms. The retrieved results, which include brief descriptions of the TCR-antigen pairs, are presented in a new table. Clicking on ‘ID’ in the first column opens a results page with more detailed interaction features, including inferred sequences, structures and binding affinities for the matching term. TCR annotation such as CDR3α, CDR3β, V(D)J recombination, species, sequencing methods, and antigen annotations involving peptide sequences, MHC allele, gene of epitope, species of epitope, as well as interactive information and experimental details such as affinity, identification and verification methods, sample and pathology are also presented. More importantly, the structure of the TCR-antigen complexes is linked with the interactive information described above.

**Figure 2.**
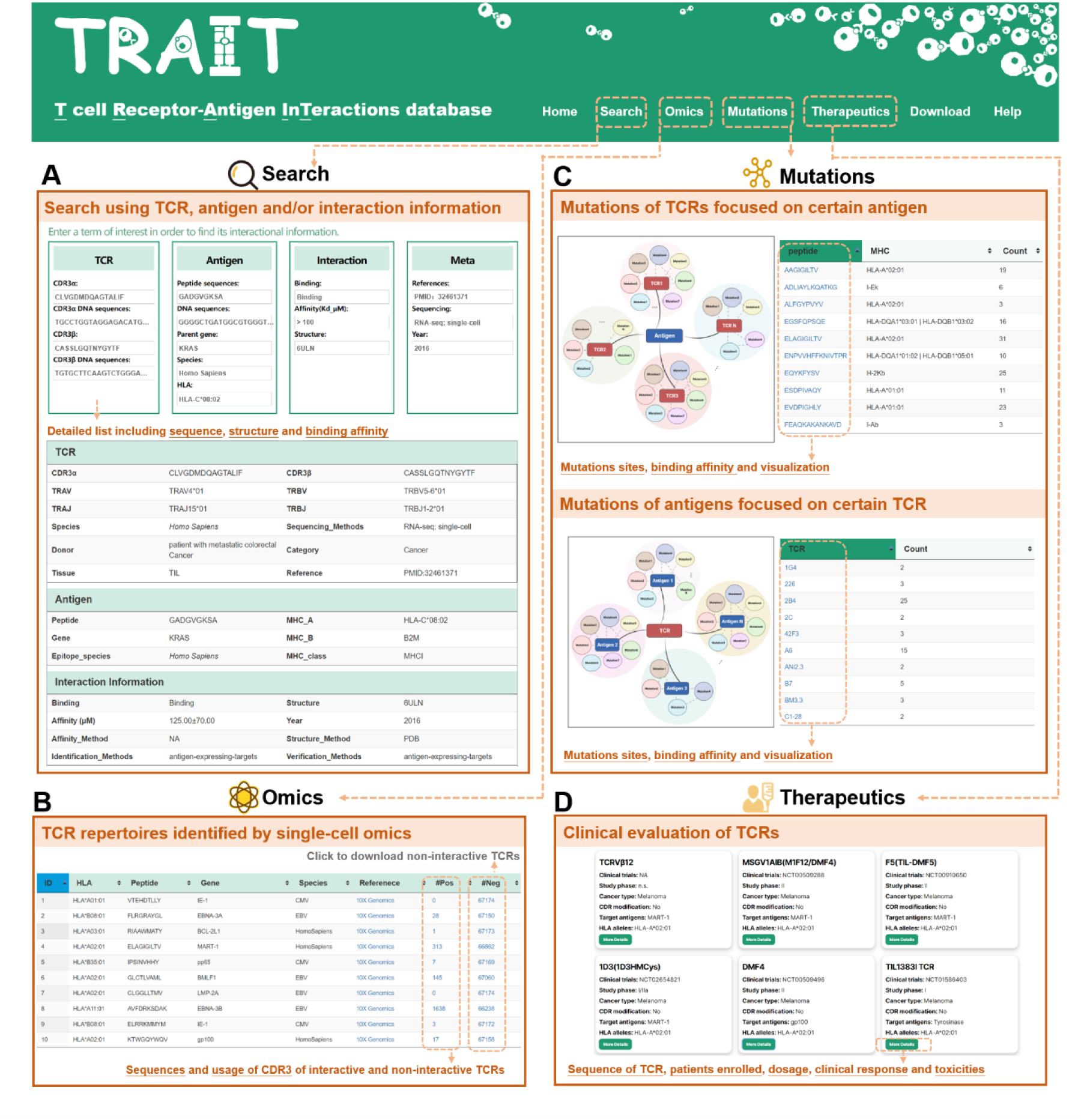
Schematic view of the database interface. **A.** The search function of TRAIT. Users can browse the detailed information including sequences, structures and binding affinities of TCR-antigen pairs. **B.** The ‘Omics’ page of TRAIT. Validated antigen-specific TCRs including interactive and non-interactive TCRs by single-cell sequencing were displayed on the page. **C.** The ‘Mutations’ page of TRAIT. Mutants of TCRs and antigens were classified and linked with their binding affinities on the page. Users are enabled to visually perceive changes on binding affinities between wildtype and mutants. **D.** The ‘Therapeutics’ page of TRAIT. The page offers detailed clinical information of TCRs on clinical trials.

The ‘Omics’ page offers a query of TCR-antigen pairs, including interactive and non-interactive pairs verified by single-cell sequencing. Clicking on ‘#pos’ in the sixth column opens a separate page for the specific antigen, showcasing detailed information about interactive TCRs and visualizations of all TCRs involved with the antigen. Users can download non-interactive TCRs for certain antigens by clicking the button ‘#neg’. The ‘Mutations’ page provides existing experimentally verified mutations of TCR-antigen pairs and their corresponding binding capacities. Clicking the TCR or antigen of interest opens a separate page for the list of all mutants and visualization of the affinity and sequence alignment. The ‘Therapeutics’ page provides a detailed overview of the pharmaceutical information for each studied TCR in clinical status. Users may click on the ‘Detail’ button to access more information, including sequences of TCR-antigen pairs, targeted disease indications, clinical status, and various clinical response data such as the number of enrolled patients, administration dosage, ORRs, and toxicities related to the studied TCR-T cells. Users can download data and obtain tutorials from the ‘Download’ and ‘Help’ pages, respectively.

## Database application

### Systematic description of TCR-antigen interactions from multiple perspectives

The engagement of TCRs with antigens constitutes one of the most complex interactions in biological systems, further complicated by the intricate mechanisms of signaling and selection [29–32]. Various experimental approaches have attempted to assess the importance of TCR-antigen interactions and correlate corresponding affinities with the level of T-cell activity mediated by antigens [10,33–36]. TRAIT presents a detailed panorama of specificity and affinity, focusing on their interplay with the structural aspects of TCRs and the interfaces they form with pMHC. As shown in Figure 2A, records of interactive TCR-antigen pairs describing antigen-specific TCR sequences are the primary source of TRAIT. A detailed annotation of the interactive pairs, including sequences of CDR3 regions, germline gene usage, sequencing methods of TCR, MHC alleles, epitope species, epitope gene of antigen, and pathologic conditions, was extracted from experimental publications. Taking mutant KRAS protein as an example, TRAIT offered comprehensive knowledge of KRAS epitopes and cognate TCRs (Table S1). A total of 35 interactive entries with 6 structures and 6 affinity data were retrieved after searching with the item ‘Parent gene: KRAS’ on the ‘Search’ page, which helps with a comprehensive understanding of KRAS and RAS drug discovery. We concluded by emphasizing how the relationships between specificity, affinity and structure might be exploited to help provide a deeper understanding of the features of antigen recognition in T-cell immunity.

### Single-cell omics-based profiling of interactions between TCRs and antigens

Single-cell sequencing provides a powerful technique to ascertain the TCR sequences of individual T-cells [37–41]. As shown in Figure 2B, TRAIT offers an omnidirectional single-cell omics map of TCRs focused on certain antigens. Detailed information on TCR-antigen pairs, including sequences of CDR3 regions and germline gene usage are shown in a separate table. A comprehensive illustration of CDR3 sequence motifs, including interactive and non-interactive pairs, is provided and can be interactively accessed online. Highly reliable and enormous TCR-antigen pairs with both interactive and non-interactive pairs will further facilitate the development of bioinformatics algorithms [42–45] and enable sample-sample comparisons within different donors and cancer types. By incorporating non-interactive pairs, a comprehensive portrayal of the binding characteristics of TCR-antigen duos is illuminated in a more comprehensive perspective.

Taking the peptide ‘GILGFVFTL’ presented by HLA-A*02:01 as an example, it has 1,368 entries of interactive and 6,5924 non-interactive TCR records from four different donors. As shown in **Figure 3**, the amino acid residues of CDR3 are relatively different between the interactive and non-interactive TCRs related to GILGFVFTL-HLA-A*02:01. CDR3 loops of most TCRs start with cysteine and alanine and end with phenylalanine in either interactive or non-interactive CDR3 motifs. Despite the subtle similarities, notable differences between interactive and non-interactive CDR3 motifs are observed. As shown in CDR3α of interactive TCRs, glycine and asparagine are preferred separately at positions 8 and 9, and phenylalanine shows biases at position 12. Preferences of amino acids in CDR3β are more significant. Interestingly, no significant amino acid preferences are observed for CDR3α and CDR3β for non-interactive TCRs except for the subtle enrichment of phenylalanine at positions 13 and 14.

**Figure 3.**
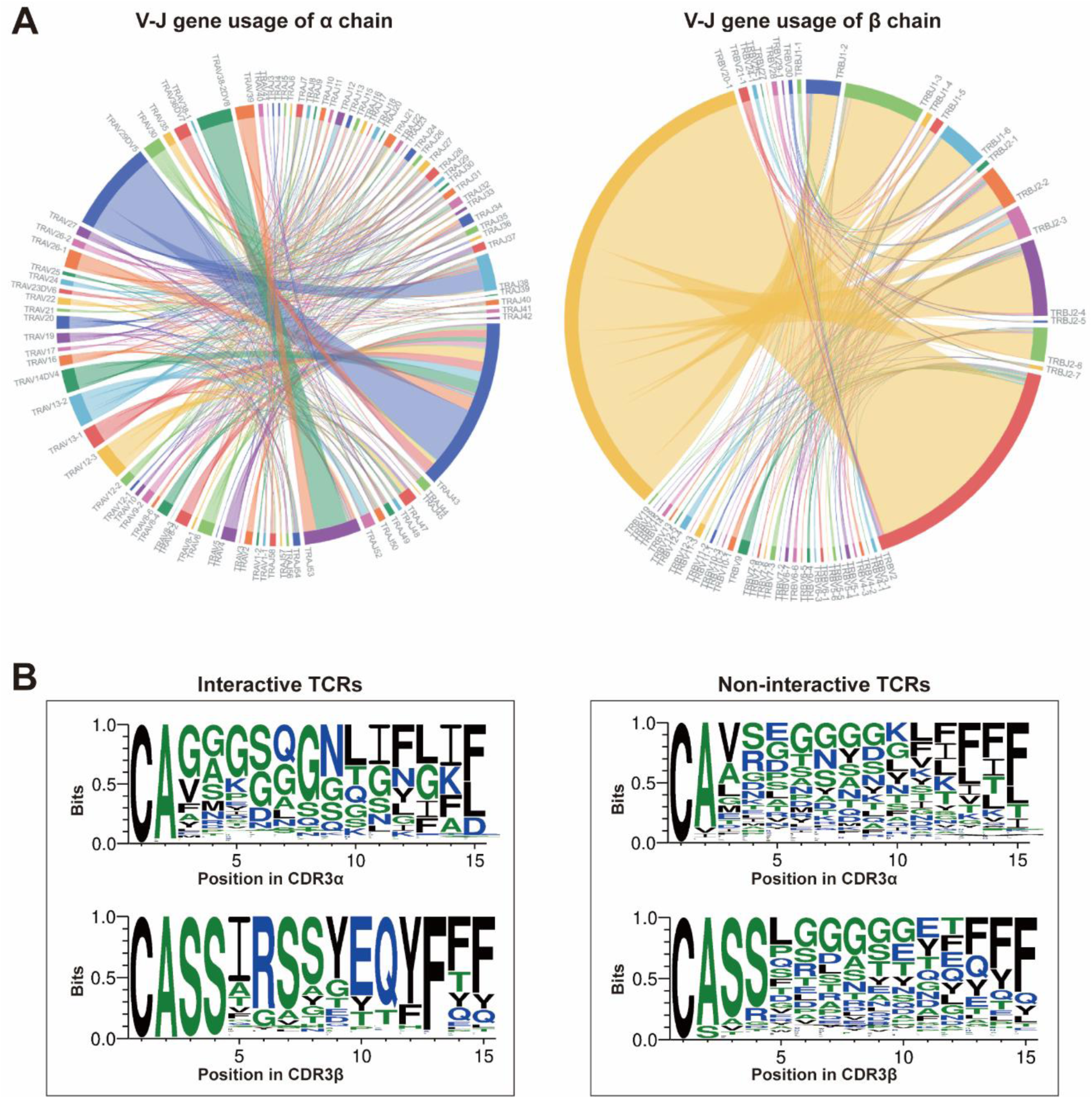
Features of TCRs related to GILGFVFTL-HLA-A*0201 on the ‘Omics’ page. **A.** V-J gene usage of interactive TCRs. The usage of genes more than five was used for visualization. **B.** Comparison of the CDR3 motif between interactive and non-interactive TCRs. The visualized figures for V-J genes on the web pages were generated using the d3.js tool. Seqlogo depicting interactive and non-interactive TCR-antigen pairs was generated using the weblogo3 tool.

### Thorough investigation of interactions between TCR mutations and cognate antigens

The manufacturing of high-affinity TCRs has been a core strategy to improve the efficacy of adoptive cell transfer therapies [46–50]. A panorama of affinities for TCR mutations and antigens was collected and systematically described in TRAIT (Figure 2C). All mutants of antigen-specific TCRs that have been experimentally verified are linked with measured affinities, and detailed information on mutations can be further consulted. Users may click SLLMWITQC-HLA-A*02:01 as an example, and 51 entries, including 42 mutation records of TCR named 1G4 and 9 mutation records of TCR named BC1, are provided in the resulting list (Table S2). Mutants of 1G4 had affinities ranging from 0.048 nM to >240 μM in four publications, while mutants of BC1 had an affinity of 0.015-21.4 μM with SLLMWITQC-HLA-A*02:01. Amino acid mutations in CDR regions were bolded and marked in red for easy track. Additionally, TRAIT enabled the visualization of the entire panorama through a scatter plot and a circular plot between mutations of TCRs and corresponding affinities with SLLMWITQC-HLA-A*02:01 (**Figure 4**A, Figure 4B). Different types of mutations (including single-point mutations, two-point mutations, and multi-point mutations) are indicated in the diagram using different colored circles, with the detailed amino acid residues interactively viewed by placing the mouse on any of the circles. Consequently, mutations of TCRs with enhanced or weakened binding capacities were intuitively viewed. In summary, TRAIT is unique in providing a panorama that enables a full visualization of the mutation affinities of TCRs and is valuable for the manufacturing of high-affinity TCR profiles for research interest. Moreover, mammalian cells [51,52], yeast [53,54], and bacteriophage display [55,56] represent additional powerful, high-throughput methods capable of creating TCRs with a wide range of binding affinities, thereby providing a continuous data source for TRAIT.

**Figure 4.**
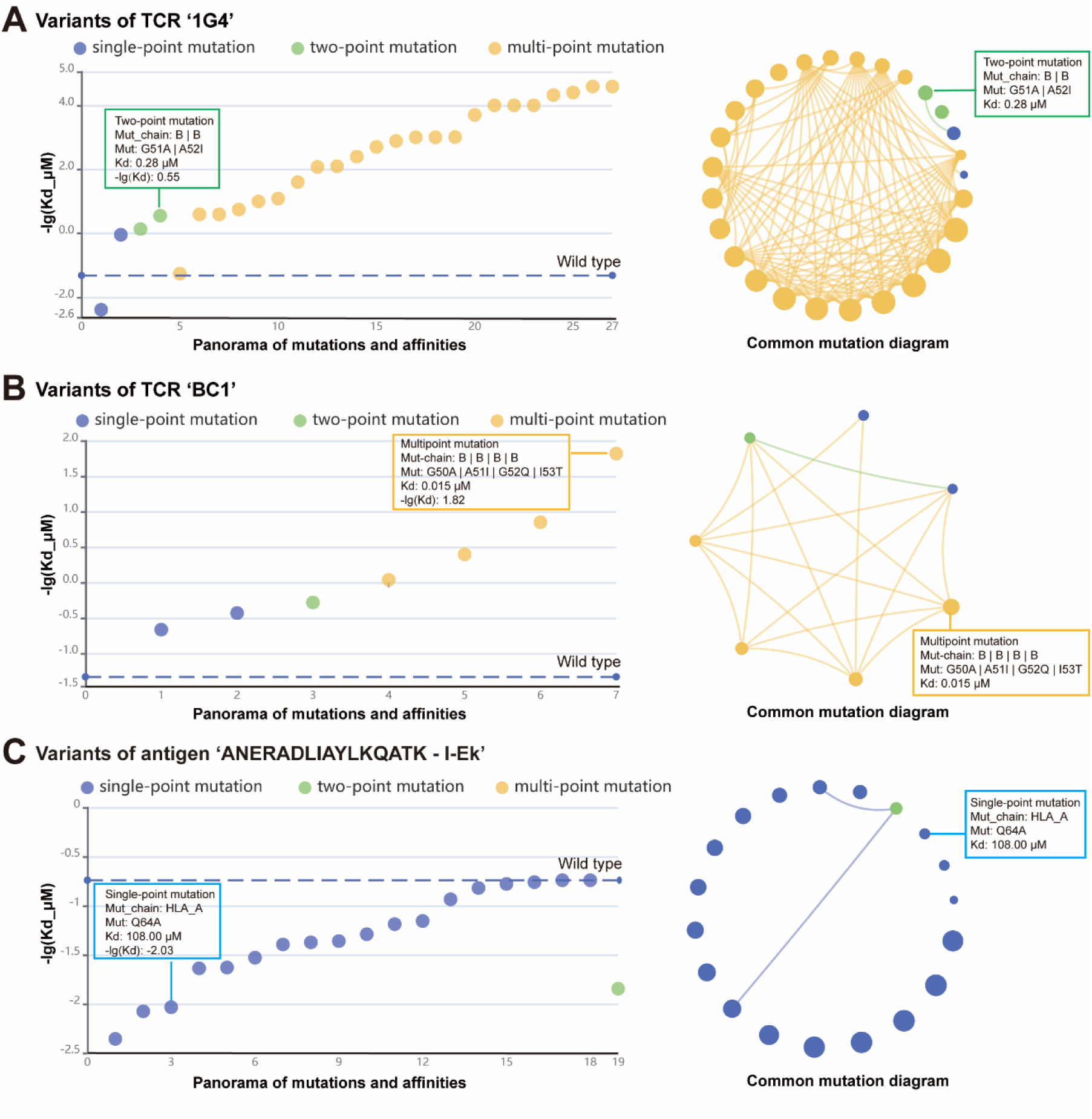
Description and visualization of binding affinity between mutations of TCR and antigens. **A**. Scatter plot and circular plot illustrating interactive capacities between mutations of TCR ‘1G4’ targeting ‘SLLMWITQC-HLA-A*02:01’. **B.** Scatter plot and circular plot illustrating interactive capacities between mutations of TCR ‘BC1’ targeting ‘SLLMWITQC-HLA-A*02:01’. **C**. Scatter plot and circular plot illustrating interactive capacities between antigen mutations and cognate TCR ‘2B4’. Mutants with shared amino acid mutation sites were visualized by linking them using lines. Larger diameters of a mutant indicate higher affinities between TCRs and their corresponding antigens. Echarts was used to visualize the relationship between mutations and affinities in online TRAIT.

### Intensive explorations of interactions between mutations of antigens and corresponding TCRs

Investigating interactions between antigens and particular TCRs will lead to a more profound understanding on immunogenicity of antigens and cross-reactivity of TCRs. TRAIT provides the binding capacities of specific TCRs with antigens or antigen mutants. As shown in Figure 2C, mutations of antigens with the capacity interacting with TCRs are summarized in the table. Detailed information can be further queried by clicking the name of TCRs and a scatter plot and a circular plot between mutants of antigens and binding capacities with certain TCRs were drawn to directly show changes in affinities to different antigens for the same TCR. Users may click 2B4 as an example, and 25 entries, including 3 wild-type record, 16 mutation records of HLA and 6 mutation records of peptides, are displayed in the search list. The scatter plot and circular plot between mutants of antigens and affinities with 2B4 indicated that neither a mutant of HLA nor peptide can result in higher affinities with TCR 2B4 (Figure 4C). Linking binding affinities to mutants of antigens enables us to delve deeper into the interactions between antigens and TCRs and help us trace peptides that can better stimulate T-cell immunity and promote vaccine development.

### Clinical usage

TCR-driven therapies are introducing a new era in precision oncology treatments. As a promising therapeutic alternative, TCR-T cells can target epitopes derived from both surface and intracellular for tumor recognition [57]. Natural (wild-type) TCRs identified from tumor-infiltrating lymphocytes, circulating T cells in patients, healthy donors, or humanized mice are the main sources of TCR candidates [58]. Nonetheless, natural TCRs typically exhibit lower affinities due to thymic selection, which restricts their capacity to recognize antigens with low densities on tumor cells. To address this issue, a common strategy is to enhance the TCR’s affinity for its targets [59]. Several TCRs without CDR modification have demonstrated encouraging outcomes in clinical studies for various solid tumors, as shown detailed in Table S3. Notably, an affinity-enhanced TCR-1G4 (targeted NY-ESO-1) achieved a 44% ORR among patients with metastatic melanoma [60–62], a 45-67% ORR in individuals with metastatic synovial sarcoma, and a 80% ORR in patients with multiple myeloma without apparent side effects [59]. Another phase 1 clinical trial indicated that TCR-T cells engineered for high affinity to MAGE-A3 antigen led to a 56% ORR in individuals with melanoma, synovial sarcoma and esophageal cancer patients [63]. In this context, the information on both natural and modified TCRs provided in TRAIT is useful for the discovery of therapeutic TCR candidates for TCR-T therapies and other immunotherapeutics.

## Conclusion and future direction

In this study, TRAIT was introduced to provide a comprehensive TCR-antigen interaction landscape. It is unique in systematically providing TCR-antigen pairs by integrating sequences, structures, binding affinities, mutations, and clinical evaluations. More importantly, a correlation between affinity and amino acid residues was initially introduced in TRAIT to more deeply explore the interaction rules and characteristics of TCRs and antigens. Moreover, it supplies a versatile visualization platform for TCR-antigen interactions, along with an intuitive web interface for convenient data retrieval. Consequently, TRAIT will serve as an invaluable resource for researchers in both experimental and computational biologists when delving into T-cell immunology and translational medicine of immunotherapies.

Immunotherapies involving the interaction of TCRs and antigens have emerged as pivotal therapeutic strategies in oncology, with an exponentially increase expected in the future for novel TCR-antigen pairs derived from patients with different diseases [64,65]. In this regard, TRAIT will be regularly updated to provide timely data and improved web pages. Furthermore, additional annotation information encompassing detailed corresponding diseases and a focus on neoantigens, along with practical analysis tools, will be incorporated to keep pace with the rapidly evolving research landscape. These enhancements are anticipated to significantly augment the application’s efficiency within the research community. Overall, TRAIT is poised to serve as a valuable resource facilitating a comprehensive understanding of molecular mechanisms underlying antigen recognition in T-cell immunology, thereby holding immense implications for the future advancements in immunotherapeutic interventions.

## Data availability

TRAIT is freely available at https://pgx.zju.edu.cn/traitdb to all users without any login or registration restrictions.

## CRediT author statement

**Mengmeng Wei:** Methodology, Data curation, Formal analysis, Investigation, Writing - original draft, Writing - review & editing. **Jingcheng Wu:** Methodology, Data curation, Software, Investigation. **Shengzuo Bai:** Software, Investigation, Visualization, Writing - review & editing. **Yuxuan Zhou:** Data curation, Investigation. **Yichang Chen:** Investigation, Visualization. **Xue Zhang**: Visualization. **Wenyi Zhao:** Methodology, Visualization. **Ying Chi:** Investigation. **Gang Pan:** Supervision. **Feng Zhu:** Conceptualization, Methodology. **Shuqing Chen:** Conceptualization, Funding acquisition. **Zhan Zhou:** Conceptualization, Funding acquisition, Supervision, Writing - review & editing. All authors have read and approved the final manuscript.

## Acknowledgments

This work was supported by the National Key Research & Developmental Program of China (Grant No. 2022YFA1305800), the National Natural Science Foundation of China (Grant No. U20A20409 and 31971371), the Zhejiang Provincial Natural Science Foundation of China (Grant No. LDT23H19011H19 and LQ24H300005), and the Huadong Medicine Joint Funds of the Zhejiang Provincial Natural Science Foundation of China (Grant No. LHDMZ22H300002). We thank the Information Technology Center and State Key Lab of CAD&CG, the Innovation Institute for Artificial Intelligence in Medicine, Zhejiang University, and Alibaba Cloud for the support of computing resources. We gratefully acknowledge all data contributors and their originating laboratories, on which the paper is based.

## Supplementary material

**Table S1 Result list by searching with ‘Parent gene: KRAS’ in TRAIT**

**Table S2 List of TCR mutations targeting SLLMWITQC-HLA-A*02:01**

**Table S3 List of representative TCRs on clinical trials**

